# Curcumin and turmeric extract inhibit SARS-CoV-2 pseudovirus cell entry and spike-mediated cell fusion

**DOI:** 10.1101/2023.09.28.560070

**Authors:** Endah Puji Septisetyani, Dinda Lestari, Komang Alit Paramitasari, Pekik Wiji Prasetyaningrum, Ria Fajarwati Kastian, Khairul Anam, Adi Santoso, Kartini Eriani

## Abstract

Turmeric extract (TE) with curcumin as its main active ingredient has been studied as a potential COVID-19 therapeutic. Curcumin has been studied in silico and in vitro against a naive SARS-CoV-2 virus, yet little is known about TE’s impact on SARS-CoV-2 infection. Moreover, no study reveals the potential of both curcumin and TE on the inhibition of SARS-CoV-2 cell-to-cell transmission. Here, we investigated the effects of both curcumin and TE on inhibiting SARS-CoV-2 entry and cell-to-cell transmission using pseudovirus (PSV) and syncytia models. We performed a PSV entry assay in 293T or 293 cells expressing hACE2. The cells were pretreated with curcumin or TE and then treated with PSV with or without the test samples. Next, we carried out syncytia assay by co-transfecting 293T cells with plasmids encoding spike, hACE2, and TMPRSS2 to be treated with the test samples. The results showed that in PSV entry assay on 293T/hACE/TMPRSS2 cells, both curcumin and TE inhibited PSV entry at concentrations of 1 µM and 10 µM for curcumin and 1 µg/ml and 10 µg/ml for TE. Moreover, both curcumin and TE reduced syncytia formation compared to control cells. Our study shows that TE and curcumin are potential inhibitors of SARS-CoV-2 infection at entry points, either by direct or indirect infection models.

**GRAPHICAL ABSTRACT:** 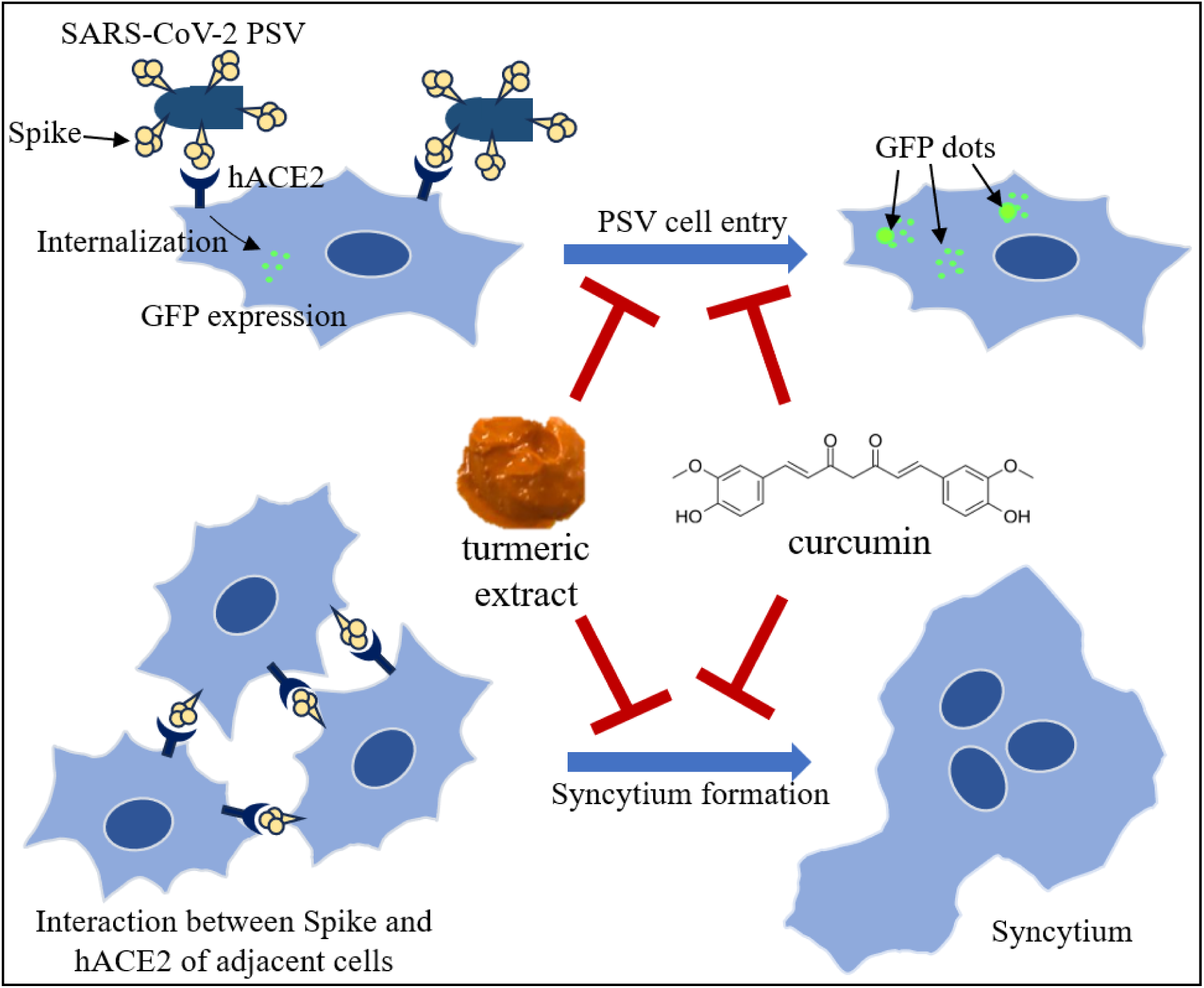

## 1. INTRODUCTION

SARS-CoV-2 can infect the target cells by direct viral infection and cell-to-cell transmission.^1,2)^ The first mode of viral infection has been widely explored by studying cell and organ tropisms. SARS-CoV-2 infection toward various cell types has been performed, which includes studies using pulmonary cancer cells Calu-3, kidney cells 293FT, liver cells Huh7, colon cancer cells CaCO2, and Vero.^3,4,5)^ In addition, organ tropisms have been studied using post-mortem tissue samples from COVID-19 patients and organoid models.^6)^ Based on these studies, SARS-CoV-2 tropisms are strongly related to hACE2 expression as its target receptor and TMPRSS2 protease, which enhances the viral entry process. SARS-CoV-2 infection involves the interaction of spike glycoprotein with hACE2, with or without spike priming by TMPRSS2, to facilitate cell entry through endocytosis or membrane fusion mechanisms.^1,7)^

In addition, SARS-CoV-2 can be transmitted through a cell-to-cell mechanism, which induces cell-to-cell fusion and syncytia formation.^8)^ Syncytia has been found in the lungs of patients infected with SARS-CoV-2.^9)^ SARS-CoV-2 infection induces spike glycoprotein expression. Eventually, as a transmembrane protein, spike protein will be transported to the plasma membrane with the receptor binding domain (RBD) at the extracellular position, ready for receptor binding. Similar to the virally integrated spike as the outer protein, cellular spike protein can also bind to the adjacent cells’ hACE2 receptor, which further induces cell-to-cell fusion and finally forms a multinucleated giant cell or syncytium.^8,9)^ Syncytia formation has been implicated with the severity of COVID-19 prognosis by several mechanisms, including immune cell phagocytosis, antibody evasion, and induction of syncytial cell death.^10)^

Due to the availability of registered COVID-19 drugs and the continuous emergence of daily COVID-19 cases, alternative treatments that can be easier to obtain for preventing and treating future COVID-19 cases are essential to be developed. Turmeric (Curcuma longa L.) is a WHO-selected medicinal plant growing in tropical areas.^11)^ Turmeric rhizome has been processed into turmeric decoction and turmeric extract, and some have been developed as a standardized medicinal extract. Turmeric extract contains curcuminoid active compounds, with curcumin as its main component.^12)^ Compared to the control group, Clinical studies have demonstrated the effect of nano-encapsulated curcumin in reducing clinical manifestations of COVID-19, such as fever, cough, and dyspnea.^13,14)^

In silico analysis showed that curcumin has a high affinity for binding to the spike glycoprotein through the formation of six hydrogen bonds,^15)^ thus, curcumin has the potential to prevent the binding of the viral spike protein to the ACE2 receptor and inhibit the initiation of the host cell infection process. Moreover, curcumin forms four hydrophobic interactions via hydrogen bonds with TMPRSS2, which may inhibit cellular entry through the cell fusion mechanism.^16)^

A plaque assay study in Vero cells has shown that curcumin inhibits SARS-CoV-2 infection in pre- and post-treatment of the D614 strain and Delta variant.^17)^ A study by Bormann *et al.* shows that turmeric root juice and curcumin showed inhibition of SARS-CoV-2 infection in Vero and Calu-3 cells determined by plaque assay and in-cell ELISA to detect the signal of SARS-CoV-2 nucleocapsid protein (N).^18)^ Nonetheless, the effect of turmeric extract (TE) on SARS-CoV-2 infection is less studied than curcumin. Moreover, there is no report about the effects of both curcumin and TE on SARS-CoV-2 cell-to-cell transmission. In the present study, we examined the effects of TE and curcumin on inhibiting SARS-CoV-2 infection using the pseudovirus (PSV) and syncytia models for targeting viral entry points and cell-to-cell transmission.

## 2. MATERIALS AND METHODS

### 2.1. Cell culture and reagents

The 293T (ECACC 12022001), CHO-K1 (ECACC 85051005), and 293 (ECACC 85120602) cell lines are collections of the Research Center for Genetic Engineering, National Research and Innovation Agency (BRIN, Indonesia). 293T cells were cultured in High-Glucose Dulbecco’s-modified Eagle’s medium (Gibco, Billings USA) supplemented with 10% (v/v) of heat-inactivated fetal bovine serum (Sigma-Aldrich, St. Louis USA) and antibiotics (100 μg/ml streptomycin and 100 U/ml penicillin). CHO-K1 cells were cultured in F-12 medium (Sigma-Aldrich, St. Louis USA) supplemented with 10% (v/v) of heat-inactivated fetal bovine serum and antibiotics (100 μg/ml streptomycin and 100 U/ml penicillin). 293 cells were cultured in MEM medium (Sigma-Aldrich, St. Louis USA) supplemented with 10% (v/v) of heat-inactivated fetal bovine serum, 1% NEAA (Gibco, Billings USA), and antibiotics (100 μg/ml streptomycin and 100 U/ml penicillin). Cells were grown inside a 37 °C tissue culture incubator at 5% CO_2_. The pcDNA3.1-SARS2-Spike (a gift from Fang Li; Addgene plasmid #145032),^19)^ pcDNA3.1-hACE (a gift from Fang Li; Addgene plasmid #145033),^19)^ and TMPRSS2 (a gift from Roger Reeves; Addgene #53887) plasmids were obtained from Addgene.^20)^ Cells were transfected with expression vectors using polyethylenimine (PEI MAX® - Transfection Grade Linear Polyethylenimine Hydrochloride (MW 40,000), Polysiences).

### 2.2. Turmeric extract and curcumin preparation

Turmeric extract (TE) was purchased as soft capsules (Natural Sari Kunyit, POM TR 192333771) from a GMP-certified manufacturer (PT. Industri Jamu dan Farmasi Sido Muncul Tbk., Semarang, Indonesia). Each capsule was standardized to contain 350 mg of concentrated liquid extract, equal to 100 mg of curcuminoids. The TE was dissolved in DMSO at 50 mg/mL, and the stock was stored at −20 °C. The working solution was freshly prepared by serially diluting the stock in culture media. Curcumin was obtained from Sigma (Cat. No. C1386-10G, Lot#SLBD0850V) (Sigma-Aldrich, St. Louis, USA), reconstituted in DMSO at 50 mM, and maintained as the stock at −20 °C. For every treatment involving curcumin, the working solution was made at 10 and 100 µM concentrations by diluting the stock with complete culture media with a maximum final concentration of DMSO less than 1%.

### 2.3. MTT assay

MTT cell viability assay was carried out in CHO-K1 and 293T cells. One day before the MTT assay, 8×10^4^ 293T or 7×10^4^ CHO-K1 cells per ml were seeded onto a 96-well plate. The next day, the cells were treated and incubated with culture media containing various concentrations of TE or curcumin for about 24 h. A set of wells containing cells without treatment and another were prepared with medium only for background subtraction (blank). Following incubation, the culture media containing TE or curcumin were discarded, and cells were washed with 1X PBS. The cells were then incubated for 2 hours in 0.5 mg/mL MTT solution (3-(4,-5-dimethylthiazo-2-yl)-2,5-diphenyltetrazolium bromide). At the end of incubation, MTT reagents were discarded, and 100 µl of DMSO were added to each well and agitated at 100 rpm for 10 minutes to ensure complete formazan crystals solubilization. Finally, the absorbance of each well was recorded at 570 nm wavelength. Cell viability was calculated according to the formula: (OD_treated_-OD_blank_)/(OD_untreated_-OD_blank_) X 100%

### 2.4. Western blot

Cell lysates were prepared using ice-cold RIPA buffer (Abcam, USA) with the addition of a protease inhibitor cocktail. The total protein concentration was determined by BCA assay (Thermo Scientific). About 10-40 µg protein was subjected to SDS-PAGE, and then the resolved protein was transferred onto an activated PVDF membrane. The membrane was then incubated in blocking buffer (5% skim milk in TBS/0.05% tween-20) followed by blotting with primary antibodies (anti-SARS-CoV-2 spike (Abcam ab275759, USA) 1:2,000 or anti-β-actin (Sigma A2228, USA) 1:4,000) for 2 h at room temperature or overnight at 4°C. After washing with TBS/T, the membrane was incubated in secondary antibodies (ALP-conjugated antibodies (Abcam ab6722), 1:4,000 or IR-Dye conjugated antibodies (LI-COR IRDye-680 RD), 1:10,000) in blocking buffer for about 2 h at room temperature or overnight at 4°C. Western blot signal was detected by incubating the membrane with 1-Step^TM^ NBT-BCIP substrate solution (Thermo Scientific 34042) or observed by a LI-COR Odyssey CLx instrument.

### 2.5. Immunofluorescence staining

The 293T cells were seeded at a density of 3×10^4^ cells/well on gelatin-coated cover glass placed inside a 24-well plate. After about 3 h incubation, the cells were transfected overnight with the pcDNA3.1-hACE2 expression vector. Transfected cells were washed with 1X PBS and fixed with 4% paraformaldehyde for 10 min. Upon fixation, cells were permeabilized with 0.2 % Triton-X, then incubated for 30 min at RT with blocking buffer (1% Bovine Serum Albumin in 1X PBS) and further incubated for at least 1 hour at RT with rabbit anti-hACE2 antibody (SAB 3500978, Sigma-Aldrich) or rabbit anti-TMPRSS2 antibody (BS-6285R, Bioss) diluted in blocking buffer (1:250). Following primary antibody staining, cells were washed three times with 1X PBS and incubated with goat anti-rabbit Alexa Fluor™ 594 or 488 secondary antibody (1:1,000) for 1 hour at RT. The secondary antibody was then washed with 1X PBS three times, and the nuclei were stained with DAPI (4′,6-diamidino-2-phenylindole)-containing mountant (Abcam Ab104139). Samples were imaged with a motorized fluorescence microscope (Olympus IX83, Tokyo, Japan).

### 2.6. Preparation of SARS-CoV-2 pseudovirus

Pseudotyping was performed in 293T cells transfected with a plasmid encoding SARS-CoV-2 spike (293T/spike). Briefly, 293T/spike cells were incubated at MOI ∼3 for 1 h with pseudotyped G*ΔG-GFP rVSV (Kerafast EH1024-PM).^21)^ Then, the medium was replaced with a fresh medium containing anti-VSV-G antibody 1:2,000 to neutralize the excess of G*ΔG-GFP rVSV and left overnight in the CO_2_ incubator. A conditioned medium (CM) containing pseudotyped spike*ΔG-GFP rVSV (SARS-CoV-2 pseudovirus) was collected and spun the next day to remove cell debris. The supernatant was aliquoted and stored at −80° C before being used for PSV entry assay.

### 2.7. Pseudovirus entry assay

PSV entry assay study was performed in the target cells of 293T transiently overexpressing hACE2 and TMPRSS2 (293T/hACE2/TMPRSS2) or 293 cells stably expressing hACE2 (293/hACE2) prepared using lentivirus system. Briefly, 4×10^5^ 293T cells were grown in each well of an 8-well chamber slide (SPL Life Sciences, Pyeongtaek, South Korea) previously coated with 2% gelatin. Upon overnight incubation, the cells were transfected with vectors harboring hACE2 and TMPRSS2 by using PEI. The next day, the medium was aspirated from the cell monolayers, and cells were subjected to pretreatment with a complete medium containing TE at 10 and 100 µg/ml or curcumin at 10 and 100 µM for 30 min. Then, the medium was removed, and the PSV was added at 1:2 ratios in 300 µl of medium containing TE or curcumin. After 16 h of viral infection, the cells were fixed and mounted. Finally, the images of GFP dots that represented the internalization of pseudovirus were acquired using a motorized fluorescence microscope (Olympus IX83). The GFP dots were counted from 8 different areas and analyzed by Fiji software (National Institute of Health).

The 293/hACE2 stable cells were generated by transduction of 293 cells using lentivirus produced in 293T cells utilizing plasmids: pCMV-dR8.2-dvpr (a gift from Bob Weinberg; Addgene #8455);^22)^ pCAGGS-G-Kan (Kerafast EH1017); and RRL.sin.cPPT.SFFV/Ace2.IRES-neo.WPRE(MT129) (a gift from Caroline Goujon; Addgene plasmid #145840).^23)^ The 293/hACE2 population were then maintained in culture medium containing G418 to obtain 293 cells that stably expressing recombinant hACE2. For the PSV entry assay using 293/hACE2, the cells were grown onto gelatin-coated 8-chamber slides and incubated overnight before PSV treatment. The GFP dots were counted from 5 different areas and analyzed by Fiji software.

### 2.8. Syncytia inhibition assay

293T cells were seeded in a 24-well plate at 1.4×105 cells/ml density in DMEM supplemented with 10% FBS. After overnight incubation, the cells were co-transfected with plasmids bearing SARS-CoV-2 spike, hACE2, and TMPRSS2 and incubated for about 6 h. Subsequently, 500 μl/well of dilutions of curcumin at 1 and 10 μM or turmeric extract at 1 μg/ml and 10 μg/ml was added to cell monolayers in duplicate and incubated for about 16 h. The following day, syncytia formation was observed using an inverted microscope (Olympus CKX53), and 10 images were acquired to represent each well. The number of syncytia was calculated with Fiji software and then sorted based on the number of nuclei using 4 categories: (i) <5 nuclei, (ii) 6-10 nuclei, (iii) 11-15 nuclei, and (iv) >15 nuclei.

### 2.9. Statistical analysis

Data were presented as mean ± SD (standard deviation) as indicated in each figure. Student’s t-test calculated statistical differences and a *p-*value <0.05 was considered significant.

## 3. RESULTS

### 3.1. Cell viability assay of curcumin and TE on CHO-K1 and 293T cells

Curcumin and TE have been known to show cytotoxic effects in various cancer cells, for instance, breast cancer cells, colorectal cancer cells, and brain cancer cells.^24)^ Also, even though curcumin and TE show less cytotoxicity against normal cells at high concentrations, the toxic effects could still be observed.^25)^ Therefore, before we investigated their antiviral activities, we evaluated the cytotoxicity of curcumin and TE on CHO-K1 and 293T cells to determine the non-toxic concentration to be used in PSV entry and syncytia fusion assays.

We applied 1, 5, 10, 25, and 50 µM curcumin serial concentrations and 1, 5, 10, 25, and 50 µg/ml TE for MTT assay in CHO-K1 cells. The results showed that after 24 hours of curcumin treatment, the cell viability was 91.7, 96.09, 92.64, 39.06, and 35.19% as the concentration increased **(Fig 1a)**. In addition, after a 24-hour treatment, CHO-K1 cell viability declined according to increased TE concentrations of 95.82, 91.08, 72.61, 41.66, and 26.27 %, respectively **(Fig 1b)**. Moreover, we tested the effect of curcumin and TE in 293T cells using serial concentrations of 1, 10, and 100 µM curcumin or 1, 10, and 100 µg/ml TE for 24 h. The results indicated that the cell viability of 293T cells after curcumin treatment was 86.52, 74.18, and 11.09 % **(Fig 1c)**. In addition, the cell viability after TE treatment was 81.52, 76.54, and 5.33 %, respectively (**Fig 1d**). Higher concentrations of tested samples significantly reduced cell viability, with the 293T cells showing more susceptibility to curcumin and TE treatment than CHO-K1 cells. Therefore, to minimize the cytotoxic effect of tested samples in further assays, we applied lower concentrations of curcumin (1 and 10 µM) and TE (1 and 10 µg/ml) and treated the cells for only about 16-18 h.

**Fig 1.**
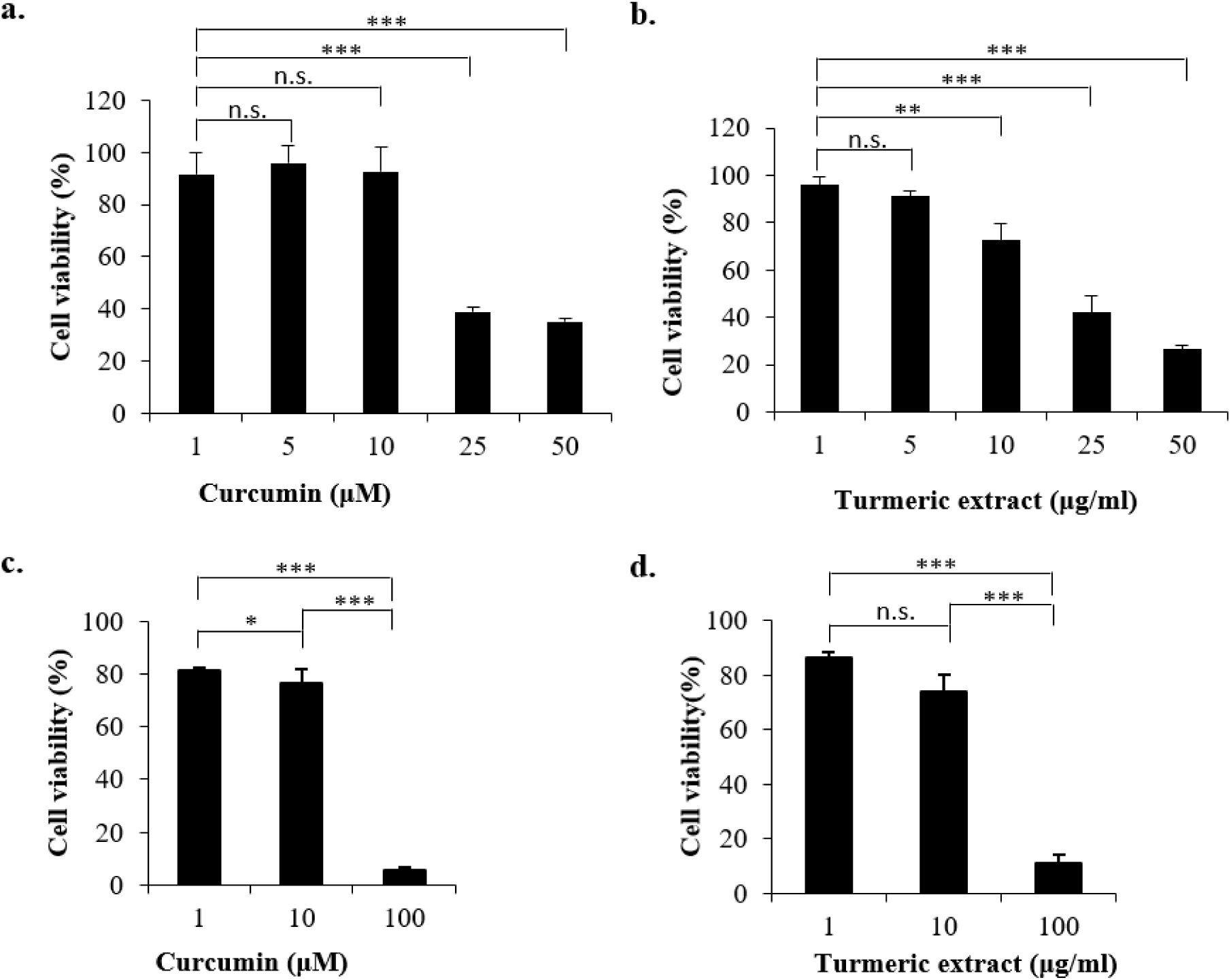
CHO-K1 (a-b) cell viability profiles and 293T cells (c-d) after curcumin and turmeric extract treatment. Cells were plated onto a 96-well plate and treated with the tested samples (n=3) for 24 h. The medium was then replaced with an MTT-containing medium and incubated for 2 h. At the end of incubation, the formazan generated was dissolved with DMSO, and the optical density was measured at 570 nm.

### 3.2. Pseudovirus entry assay

We prepared SARS-CoV-2 PSV with a VSV backbone and GFP reporter in spike-transfected 293T cells (pseudotyped spike*ΔG-GFP rVSV). Spike expression in 293T/spike cells used for pseudotyping was confirmed by Western blot. Moreover, we also detected spike in the CM obtained after pseudotyping, indicating the formation of SARS-CoV-2 pseudovirus and its release into the culture medium (**Fig 2**).

**Fig 2.**
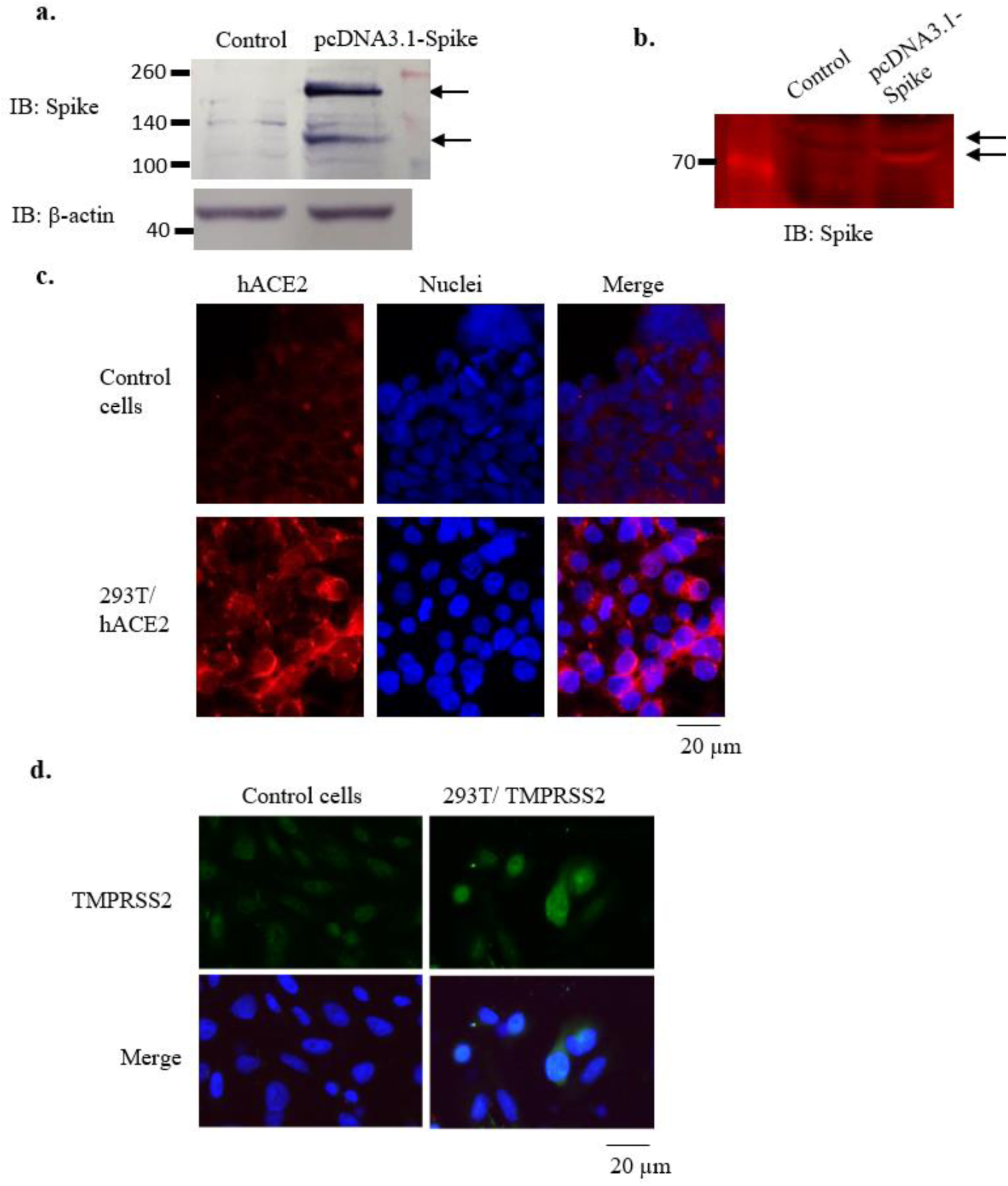
Observation of spike, hACE2, and TMPRSS2 expression. a. Detection of spike expression in 293T cells transfected with spike plasmid. b. Detection of spike in the conditioned medium, which represented SARS-CoV-2 PSV. c-d. Detection of hACE2 and TMPRSS2 expression by immunofluorescence staining.

Furthermore, we used the CM to perform a PSV entry assay in 293T overexpressing hACE2/TMPRSS2 and 293/ACE2 stable cells. The 293T/hACE2/TMPRSS2 cells were pretreated with curcumin or TE for 30 min, then treated with PSV/curcumin or PSV/TE, followed by 16-18 h incubation in a CO_2_ incubator. GFP expression indicating PSV internalization and viral genome release was observed by fluorescence microscopy with 60x magnification to expose the GFP dots **(Fig 3a)**. To confirm that GFP dots were formed within the cells, we also observed the cells in bright field mode and used DAPI to stain the nuclei. The representative images are shown in **Fig 3b**. As the results showed, nontreated cells showed a higher number of GFP dots compared to the treated cells with fluorescence focus units (FFU) 296±2. The FFU of cells treated with curcumin 1 and 10 µM were 141±8.76 and 56±6.52. Whereas FFU of cells treated with TE 1 and 10 µg/ml were 81+5.93 and 49±3.31. Cells treated with curcumin and TE significantly reduced the FFU number, especially at higher concentrations, indicating the potential inhibitory effect on SARS-CoV-2 viral cell entry (**Fig 3c**).

**Fig 3.**
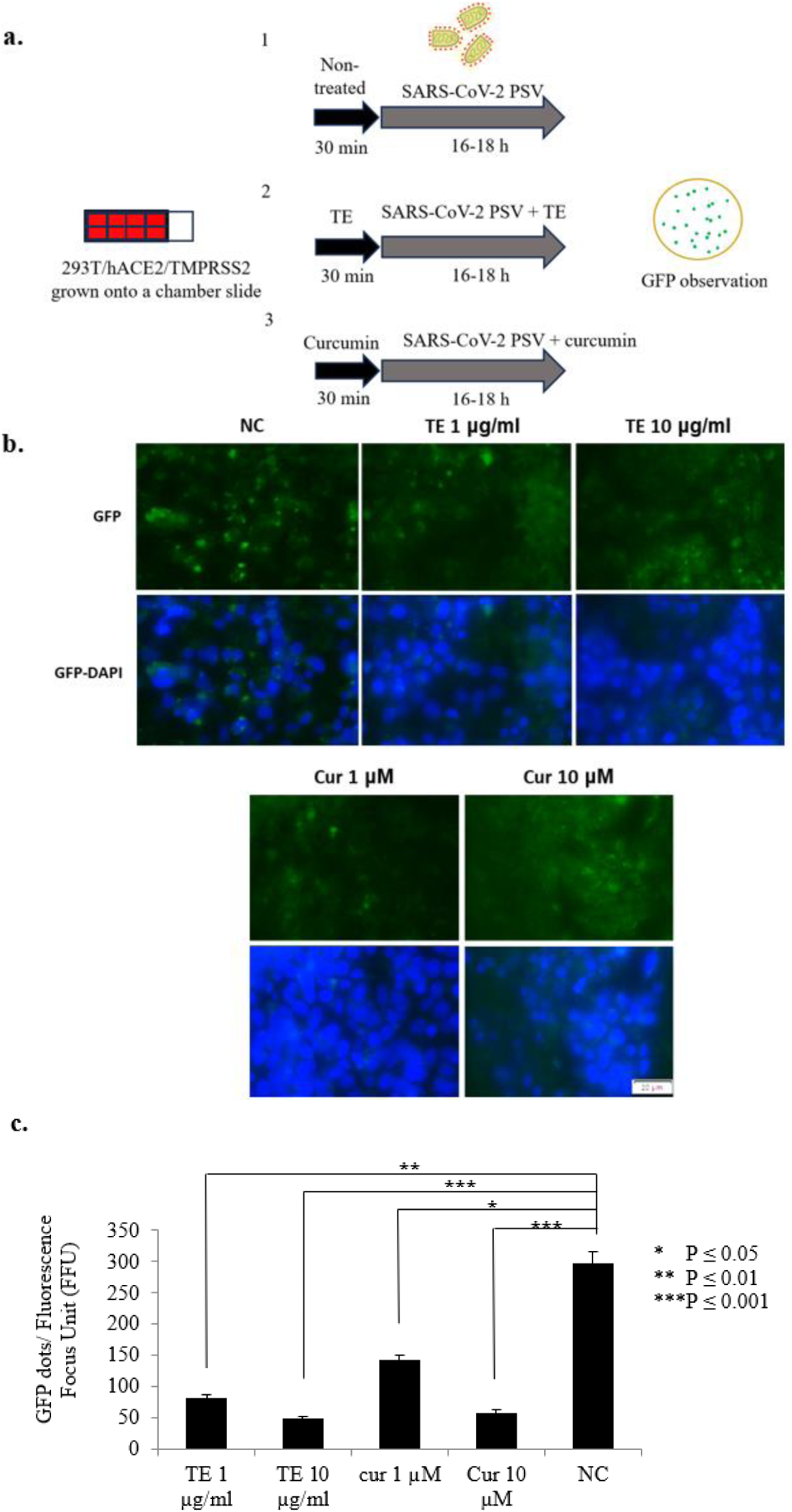
Effect of turmeric extract and curcumin on PSV entry in 293T/hACE2/TMPRSS2 cells. a. Schematic representation of PSV entry assay in 293T/hACE2/TMPRSS2 cells. b. The representative image shows GFP dots to represent PSV internalization. c. Graph representing quantification of GFP dots (n = 8 microscope fields). TE: turmeric extract; Cur: curcumin; NC: nontreated cells.

Next, we clarified the inhibitory effect of curcumin as the active compound to inhibit SARS-CoV-2 PSV entry by performing an entry assay in 293 cells that stably expressed ACE2 (293/ACE2) **(Fig 4a)**. Besides treating the 293/ACE2 cells with curcumin before PSV addition, the curcumin was combined with PSV after incubation for 30 min **(Fig 4b)**. The representative images of cells after the PSV entry assay are shown in **Fig 4c**. As a quantification result, the number of GFP dots/cell for non-treated cells was 1.45. The number of GFP dots per cell in curcumin and PSV/curcumin-treated cells was 0.99, while the number of GFP dots per cell in curcumin-treated cells was 0.887. The results indicated that curcumin reduced PSV entry, especially for curcumin pretreatment before the addition of PSV (P = 0.035) (**Fig 4d**).

**Fig 4.**
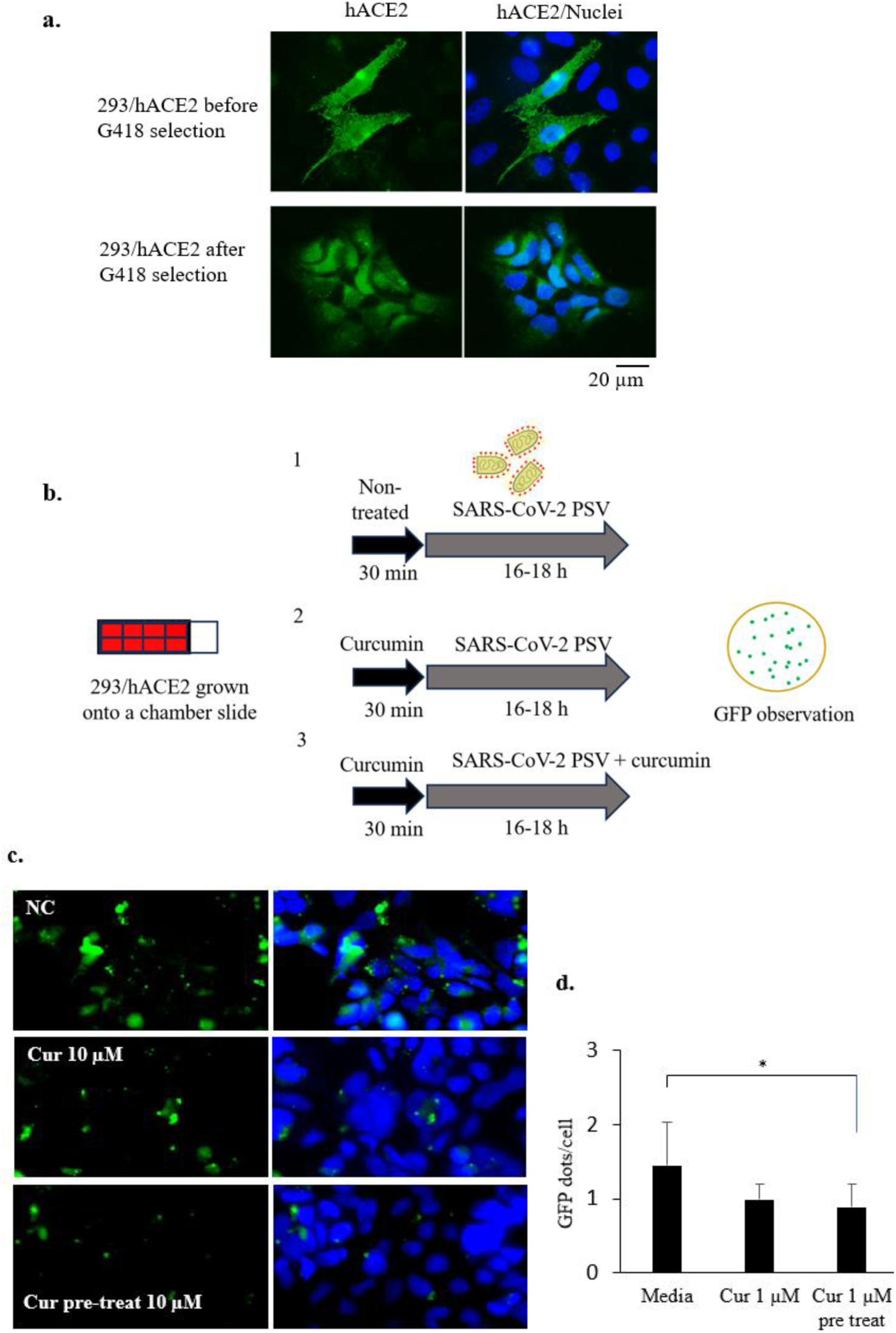
Effect of curcumin on PSV entry in 293/ACE2 cells. a. Observation of hACE2 expression in 293/hACE2 cells after transduction, before and after selection with antibiotic G418. b. Schematic representation of PSV entry assay in 293/hACE2 stable cells. c. The representative image shows GFP dots to represent PSV internalization. d. Graph representing quantification of GFP dots (n = 5 microscope fields). P<0.05 = *. NC: nontreated cells.

### 3.3. Syncytia formation assay

This assay represents the cells infected with SARS-CoV-2 and expressing its membrane protein spike, which can bind the hACE2 receptor of the adjacent cell. Instead of using the original virus, we transfected the cells directly with spike plasmid. We used 293T cells transfected with plasmids encoding SARS-CoV-2 spike, hACE2, and TMPRSS2 to perform the syncytia assay. The cells expressing spike will locate the Spike in the transmembrane region, enabling the interaction between spike with the hACE2 receptor and TMPRSS2 of the adjacent cells. As a result, the two interacting cells will form an intercellular bridge that eventually will fuse and form multinucleated cells.^26)^ The more cells interact and fuse, the more multinucleated cells or syncytia will be generated, as shown in **Fig 5a**.

**Fig 5.**
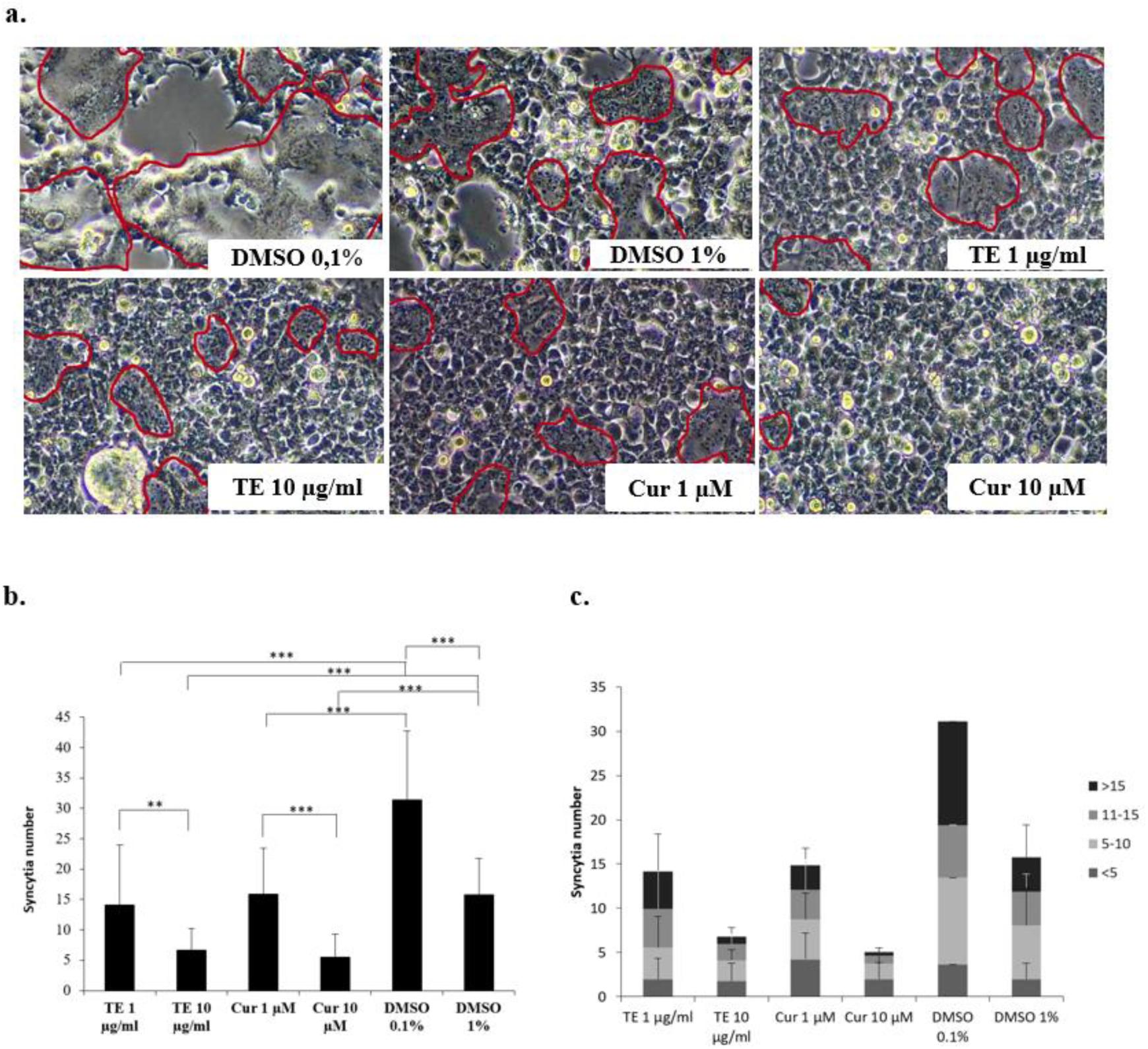
Effect of curcumin and turmeric extract on syncytia formation. a. Representative image showing the syncytia formed after curcumin, TE, and DMSO treatment. b. Graph representing quantification of syncytia number (n = 10 microscopic fields in duplicate). c. Graph representing quantification of nuclei number per categories of syncytia (n = 10 microscopic fields). Red area: syncytia. P<0.05 = *, P<0.01 = **, P<0.001 = ***, ns = not significant.

We show that 293T cells, after transfection and treatment with DMSO 0.1% and 1%, formed syncytia as much as 31.4 and 15.75 per field. In addition, the treatment of cells with curcumin and TE reduced the number of syncytia. The number of syncytia formed after cell treatment with curcumin 1 and 10 µM was 15.85 (P<0.0001 vs DMSO 0.1%) and 5.5 (P<0.0001 vs DMSO 1%), while the number of syncytia formed after cell treatment with TE 1 and 10 µg/ml was 14.05 (P<0.0001 vs DMSO 0.1%) and 6.6 (P<0.0001 vs DMSO 0.1%) (**Fig 5b**).

Furthermore, we analyzed the distribution of nuclei numbers within the syncytia and categorized them into <5, 5-10, 11-15, and <15. The syncytia with nuclei number <5 for DMSO 0.1, curcumin 1 µM, and TE 1 µg/ml were 3.65, 4.25, 2, while nuclei number 5-10 were 9.8, 4.5, 35.5, nuclei no 11-15 were 5.95, 3.3, 4.35, and nuclei number >15 were 11.75, 2.8, 4.25. The syncytia with nuclei number <5 for DMSO 1, curcumin 10 µM and TE 10 µg/ml were 2, 2, 1.8, while nuclei number 5-10 were 6.05, 1.75, 2.3, nuclei no 11-15 were 3.85, 0.9, 1.85, and nuclei no >15 were 3.85, 0.4, 0.85. These data indicated that treatment of the cells with curcumin and TE reduced the nuclei number within syncytia, representing lower fusion events than the DMSO treatment (**Fig 5c**).

## 4. DISCUSSION

Curcumin has been tested for its anti-SARS-CoV-2 activities by plaque assay in Vero cells. Using the original virus, curcumin inhibits SARS-CoV-2 infection.^17)^ However, surface receptors in Vero cells differ from human cells, and the study may not correlate well with actual events.^27)^ Thus, the antiviral study of curcumin has also been carried out using human Calu-3 cells, which endogenously express hACE2 and TMPRSS2. A study using these cells demonstrated that curcumin may inhibit SARS-CoV-2 viral replication as indicated by reduced N protein expression following viral infection.^18)^ Besides investigating curcumin’s antiviral effect, Bormann *et al*. also tested the effect of turmeric juice (water extract).^18)^ However, the antiviral study of turmeric extract (TE) has not been widely reported compared to its active compound, curcumin.

In this manuscript, we reported the SARS-CoV-2 antiviral activities of curcumin and TE, especially at the entry point, using the pseudovirus and syncytia models to represent cell-to-cell transmission. Here, we showed that curcumin and TE reduced PSV entry in 293T/hACE2/TMPRSS2 cells, in which 10 µM curcumin and 10 µg/ml TE significantly affected the number of GFP dots. The effect of TE was comparable to the curcumin effect on PSV cell entry in which 10 µg/ml TE was equal to 2.86 µg/ml curcuminoid, with the majority of curcumin content and 10 µM curcumin being equal to 3.68 µg/ml of curcumin.

From the previous studies, Marin-Palma *et al*. reported that 10 µM curcumin can inhibit SARS-CoV-2 infection in Vero E6 cells.^17)^ It is also reported that curcumin inhibits SARS-CoV-2 infection at concentrations of 3-10 µM.^28)^ Furthermore, in A549 cells, curcumin shows SARS-CoV-2 antiviral concentration of 5 µM.^29)^ We also showed that curcumin inhibited PSV entry in 293/hACE2 cells. These results corroborate curcumin effects against SARS-CoV-2 infection with our data representing curcumin inhibition at PSV cell entry point. It has been known that curcumin affects the early stages of viral replication cycles, including viral-receptor attachment, internalization, and fusion that have been studied against several types of viruses, which involve influenza, dengue, zika, chikungunya, pseudorabies, and VSV.^30,31,32)^

Moreover, curcumin and TE inhibit secondary infection via cell-to-cell transmission in a syncytia formation model mediated by SARS-CoV-2 spike expression. Cells treated with curcumin and TE showed smaller syncytia with fewer nuclei than control cells. The more cells fused to generate syncytia, the larger syncytia with more nuclei will be formed, and vice versa.

Based on in silico data, curcumin can also interact with SARS-CoV-2 spike RBD,33) hACE2,34) and TMPRSS2.^16)^ These data align with our results that curcumin inhibited PSV entry and syncytia formation. Our in vitro study using PSV and syncytia models revealed that both curcumin and TE are potential inhibitors of SARS-CoV-2 infection, especially at the entry points either by direct infection or cell-to-cell transmission mediated by spike-induced cell fusion. Curcumin can interfere with the spike-receptor binding during direct viral or intercellular transmission, hindering viral infection and cell fusion.^17)^ In addition, TE, as the crude extract that contains curcumin, also has the potential to inhibit SARS-CoV-2 infection and potentially to be developed as an independent herbal-derived product for the prevention of viral infection with curcuminoids used as identity compounds for TE standardization.

## ACKNOWLEDGEMENTS

We are grateful for the financial support from the Educational Fund Management Institution/National Research and Innovation Agency (LPDP/BRIN) (grant no. 102/FI/P-KCOVID-19.2B3/IX/2020 and RIIM No. KEP-5/LPDP/LPDP.4/2022) and Research Organization for Life Sciences and Environment, BRIN (Research Program (DIPA Rumah Program) 2022).

## AUTHORS’ CONTRIBUTION

Conceptualization: Septisetyani EP. Data curation: Lestari D, Prasetyaningrum PW, Paramitasari KA, Kastian RF, Septisetyani EP. Writing - original draft: Septisetyani EP, Lestari D, Paramitasari KA. Writing - review & editing: Septisetyani EP, Prasetyaningrum PW, Anam K, Santoso A., Eriani K.

## Supplementary

**Supplementary Figure 1.).**
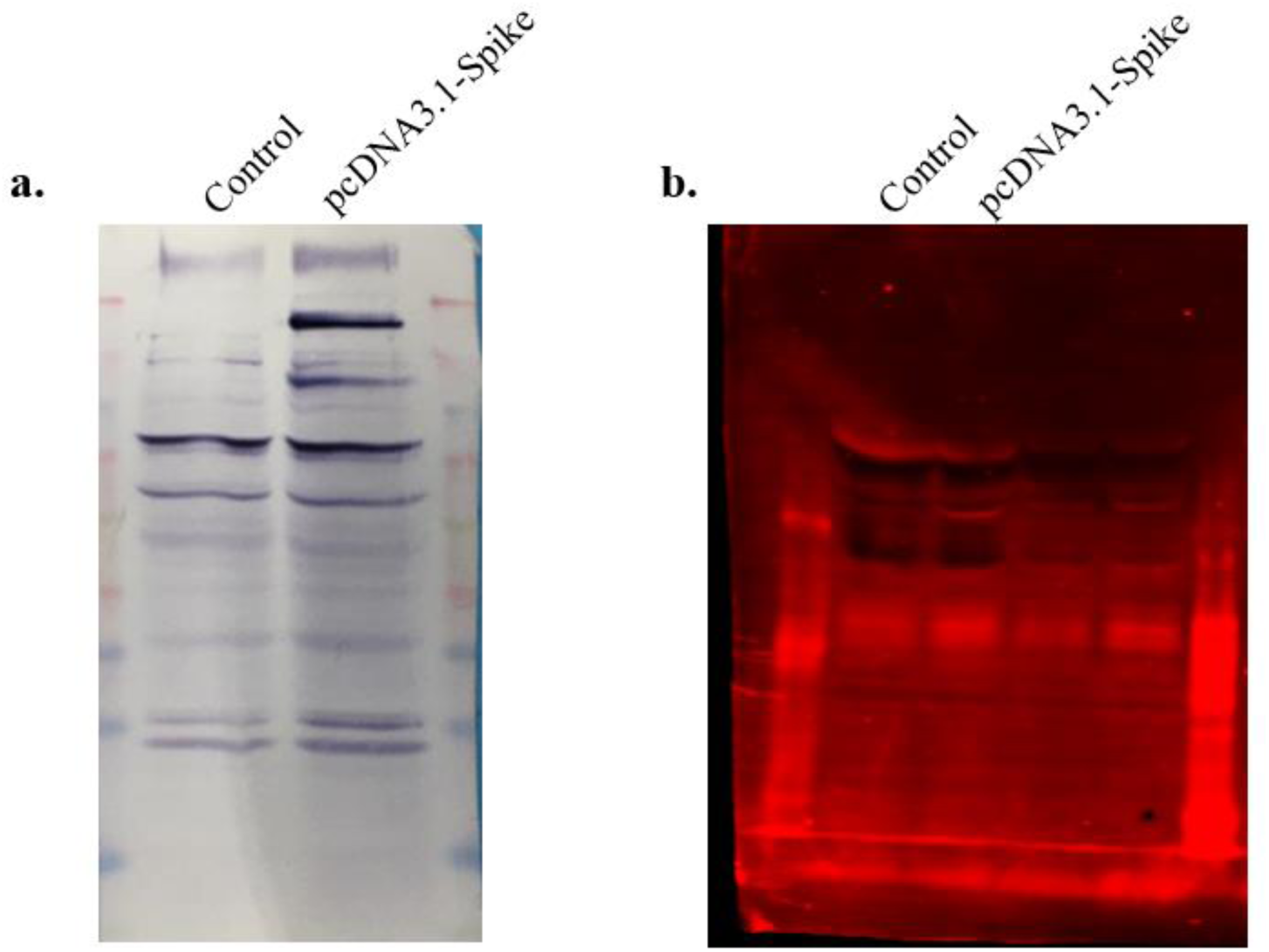
Full membrane image of spike detection by western blot.

## REFERENCES

1) Jackson CB, Farzan M, Chen B, Choe H. Mechanisms of SARS-CoV-2 entry into cells. Nat. Rev. Mol. Cell. Biol., 23, 1–18 (2021).

2) Rajah MM, Hubert M, Bishop E, Saunders N, Robinot R, Grzelak L, Planas D, Dufloo J, Gellenoncourt S, Bongers A, Zivaljic M, Planchais C, Guivel-Benhassine F, Porrot F, Mouquet H, Chakrabarti LA, Buchrieser J, Schwartz O. SARS-CoV-2 Alpha, Beta, and Delta variants display enhanced spike-mediated syncytia formation. EMBO J., 40, e108944 (2021).

3) Hoffmann M, Kleine-Weber H, Schroeder S, Krüger N, Herrler T, Erichsen S, S, Schiergens TS, Herrler G, Wu NH, Nitsche A, Müller MA, Drosten C, Pöhlmann. SARS-CoV-2 cell entry depends on ACE2 and TMPRSS2 and is blocked by a clinically proven protease inhibitor. Cell, 181, 271–280 (2020).

4) Katopodis P, Randeva HS, Spandidos DA, Saravi S, Kyrou I, Karteris E. Host cell entry mediators implicated in the cellular tropism of SARS-CoV-2, the pathophysiology of COVID-19 and the identification of microRNAs that can modulate the expression of these mediators (Review). Int. J. Mol. Med., 49, 20 (2022).

5) Saccon E, Chen X, Mikaeloff F, Rodriguez JE, Szekely L, Vinhas BS, Krishnan S, Byrareddy SN, Frisan T, Végvári Á, Mirazimi A, Neogi U, Gupta S. Cell-type-resolved quantitative proteomics map of interferon response against SARS-CoV-2. iScience, 24, 102420 (2021).

6) Peng L, Gao L, Wu X, Fan Y, Liu M, Chen J, Song J, Kong J, Dong Y, Li B, Liu A, Bao F. Lung organoids as model to study SARS-CoV-2 infection. Cells, 11, 2758 (2022).

7) Hirano T, Murakami M. COVID-19: A new virus, but a familiar receptor and cytokine release syndrome. Immunity, 52, 731–733 (2020).

8) Zheng Y, Zhou LL, Su Y, Sun Q. Cell fusion in the pathogenesis of COVID-19. Mil. Med. Res., 8, 68 (2021).

9) Lin L, Li Q, Wang Y, Shi Y. Syncytia formation during SARS-CoV-2 lung infection: a disastrous unity to eliminate lymphocytes. Cell Death Differ., 28, 2019–2021, (2021).

10) Rajah MM, Bernier A, Buchrieser J, Schwartz O. The mechanism and consequences of SARS-CoV-2 spike-mediated fusion and syncytia formation. J. Mol. Biol., 434, 167280 (2022).

11) World Health Organization. Rhizoma Curcumae Longae. WHO monographs on selected medicinal plants. Vol. 1, WHO, Geneva, pp. 115–124 (1999).

12) Sharifi-Rad J, Rayess YE, Rizk AA, Sadaka C, Zgheib R, Zam W, Sestito S, Rapposelli S, Neffe-Skocińska K, Zielińska D, Salehi B, Setzer WN, Dosoky NS, Taheri Y, El Beyrouthy M, Martorell M, Ostrander EA, Suleria HAR, Cho WC, Maroyi A, Martins N. Turmeric and its major compound curcumin on health: bioactive effects and safety profiles for food, pharmaceutical, biotechnological and medicinal applications. Front. Pharmacol., 11, 01021 (2020).

13) Tahmasebi S, El-Esawi MA, Mahmoud ZH, Timoshin A, Valizadeh H, Roshangar L, Varshoch M, Vaez A, Aslani S, Navashenaq JG, Aghebati-Maleki L, Ahmadi M. Immunomodulatory effects of nanocurcumin on Th17 cell responses in mild and severe COVID-19 patients. J. Cell Physiol., 236, 5325–5338 (2021).

14) Valizadeh H, Abdolmohammadi-Vahid S, Danshina S, Ziya GM, Ammari A, Sadeghi A, Roshangar L, Aslani S, Esmaeilzadeh A, Ghaebi M, Valizadeh S, Ahmadi M. Nano-curcumin therapy, a promising method in modulating inflammatory cytokines in COVID-19 patients. Int. Immunopharmacol., 89, 107088 (2020).

15) Maurya VK, Kumar S, Prasad AK, Bhatt MLB, Saxena SK. Structure-based drug designing for potential antiviral activity of selected natural products from Ayurveda against SARS-CoV-2 spike glycoprotein and its cellular receptor. Virusdisease, 31, 179– 193 (2020).

16) Motohashi N, Vanam A, Gollapudi R. In silico study of curcumin and folic acid as potent inhibitors of human transmembrane protease serine 2 in the treatment of COVID-19. INNOSC Theranostics and Pharmacological Sciences, 16, 3–9 (2020).

17) Marín-Palma D, Tabares-Guevara JH, Zapata-Cardona MI, Flórez-Álvarez L, Yepes LM, Rugeles MT, Zapata-Builes W, Hernandez JC, Taborda NA. Curcumin inhibits in vitro SARS-CoV-2 infection in Vero E6 cells through multiple antiviral mechanisms. *J*. Molecules, 26, 6900 (2021).

18) Bormann M, Alt M, Schipper L, van de Sand L, Le-Trilling VTK, Rink L, Heinen N, Madel RJ, Otte M, Wuensch K, Heilingloh CS, Mueller T, Dittmer U, Elsner C, Pfaender S, Trilling M, Witzke O, Krawczyk A. Turmeric root and its bioactive ingredient curcumin effectively neutralize SARS-CoV-2 in vitro. Viruses, 13,1914 (2021).

19) Shang J, Ye G, Shi K, Wan Y, Luo C, Aihara H, Geng Q, Auerbach A, Li F. Structural basis of receptor recognition by SARS-CoV-2. Nature, 581, 221–224 (2020).

20) Edie S, Zaghloul NA, Leitch CC, Klinedinst DK, Lebron J, Thole JF, McCallion AS, Katsanis N, Reeves RH. Survey of human chromosome 21 gene expression effects on early development in *Danio rerio*. G3 (Bethesda), 8, 2215–2223 (2018).

21) Whitt MA. Generation of VSV pseudotypes using recombinant ΔG-VSV for studies on virus entry, identification of entry inhibitors, and immune responses to vaccines. J. Virol. Methods, 169, 365–374 (2010).

22) Stewart SA, Dykxhoorn DM, Palliser D, Mizuno H, Yu EY, An DS, Sabatini DM, Chen IS, Hahn WC, Sharp PA, Weinberg RA, Novina CD. Lentivirus-delivered stable gene silencing by RNAi in primary cells. RNA, 9, 493–501 (2003).

23) Rebendenne A, Valadão ALC, Tauziet M, Maarifi G, Bonaventure B, McKellar J, Planès R, Nisole S, Arnaud-Arnould M, Moncorgé O, Goujon C. SARS-CoV-2 triggers an MDA-5-dependent interferon response which is unable to control replication in lung epithelial cells. J. Virol., 95, e02415–20 (2021).

24) Tomeh MA, Hadianamrei R, Zhao X. A review of curcumin and its derivatives as anticancer agents. Int. J. Mol. Sci., 20, 1033 (2019).

25) Li P, Pu S, Lin C, Liu H, Zhao H, Yang C, Guo Z, Xu S, Zhou Z. Curcumin selectively induces colon cancer cell apoptosis and S cell cycle arrest by regulates Rb/E2F/p53 pathway. J. Mol. Struct., 1263, 133180 (2022).

26) Buchrieser J, Dufloo J, Hubert M, Monel B, Planas D, Rajah MM, Planchais C, Porrot F, Guivel-Benhassine F, Van der Werf S, Casartelli N, Mouquet H, Bruel T, Schwartz O. Syncytia formation by SARS-CoV-2-infected cells. EMBO J., 39, e106267 (2020).

27) Pandamooz S, Jurek B, Meinung CP, Baharvand Z, Sahebi Shahem-Abadi A, Haerteis S, Miyan JA, Downing J, Dianatpour M, Borhani-Haghighi A, Salehi MS. Experimental models of SARS-CoV-2 infection: possible platforms to study COVID-19 pathogenesis and potential treatments. Annu. Rev. Pharmacol. Toxicol., 62, 25–53 (2022).

28) Wen CC, Kuo YH, Jan JT, Liang PH, Wang SY, Liu HG, Lee CK, Chang ST, Kuo CJ, Lee SS, Hou CC, Hsiao PW, Chien SC, Shyur LF, Yang NS. Specific plant terpenoids and lignoids possess potent antiviral activities against severe acute respiratory syndrome coronavirus. J. Med. Chem., 50, 4087–4095 (2007).

29) Goc A, Sumera W, Rath M, Niedzwiecki A. Phenolic compounds disrupt spike-mediated receptor-binding and entry of SARS-CoV-2 pseudo-virions. PLoS ONE, 16, e0253489 (2021).

30) Mounce BC, Cesaro T, Carrau L, Vallet T, Vignuzzi M. Curcumin inhibits Zika and chikungunya virus infection by inhibiting cell binding. Antivir. Res., 142, 148–57 (2017).

31) Chen DY, Shien JH, Tiley L, Chiou SS, Wang SY, Chang TJ, Lee YJ, Chan KW, Hsu WL. Curcumin inhibits influenza virus infection and haemagglutination activity. Food Chem., 119, 1346–1351 (2010).

32) Chen TY, Chen DY, Wen HW, Ou JL, Chiou SS, Chen JM, Wong ML, Hsu WL. Inhibition of enveloped viruses infectivity by curcumin. PLoS ONE, 8, e62482 (2013).

33) Shanmugarajan D, Prabitha P, Kumar BRP, Suresh B. Curcumin to inhibit binding of spike glycoprotein to ACE2 receptors: computational modelling, simulations, and ADMET studies to explore curcuminoids against novel SARS-CoV-2 targets. RSC Adv., 10, 31385–31399 (2020).

34) Subbaiyan A, Ravichandran K, Singh SV, Sankar M, Thomas P, Dhama K, Malik YS, Singh RK, Chaudhuri P. In silico molecular docking analysis targeting SARS-CoV-2 spike protein and selected herbal constituents. J. Pure Appl. Microbiol., 14, 989–998 (2020).

